# Evolutionary analyses reveal independent origins of gene repertoires and structural motifs associated to fast inactivation in calcium-selective TRPV channels

**DOI:** 10.1101/564997

**Authors:** Lisandra Flores-Aldama, Michael W. Vandewege, Kattina Zavala, Charlotte K. Colenso, Wendy Gonzalez, Sebastian E. Brauchi, Juan C. Opazo

## Abstract

Important for calcium homeostasis, TRPV5 and TRPV6 are calcium-selective channels belonging to the transient receptor potential (TRP) gene family. In this study, we investigated the evolutionary history of these channels to add an evolutionary context to the already available physiological information. Phylogenetic analyses revealed that paralogs found in mammals, sauropsids, amphibians, and chondrichthyans, are the product of independent duplication events in the ancestor of each group. Within amniotes, we identified a traceable signature of three amino acids located at the amino-terminal intracellular region (HLH domain). The signature correlates well with both the duplication events and the phenotype of fast inactivation observed in mammalian TRPV6 channels. Electrophysiological recordings and mutagenesis suggest that calcium-induced fast inactivation represents an evolutionary innovation that emerged independently after gene duplication.

## Introduction

In vertebrates, epithelial calcium absorption is a crucial physiological process needed to maintain Ca^2+^ homeostasis. Epithelia consist of a continuous layer of individual cells where calcium absorption is maintained by two routes, the transcellular and paracellular pathways (Goor et al., 2017; Hoenderop et al., 2005). Tight junctions regulate the passive passage of ions and molecules through the paracellular pathway (Hoenderop et al., 2005). Calcium permeation through this pathway constitutes a major route for Ca^2+^ absorption under normal physiological conditions (Goor et al., 2017). On the other hand, the transcellular pathway implicates controlled Ca^2+^ movement through epithelial barriers and occurs against a concentration gradient in normal conditions. It is an active and saturable process, maintained by an array of transporters, pumps, and ion channels (Goor et al., 2017).

TRPV5 and TRPV6 are calcium-selective ion channel members of the Transient Receptor Potential (TRP) gene family (Van Abel et al., 2005). These proteins, expressed at the apical membrane of Ca^2+^ transporting epithelia, serve as entry channels in transepithelial Ca^2+^ transport (Van Abel et al., 2005). In agreement with their physiological role, TRPV6 deficient mice have impaired Ca^2+^ homeostasis, evidenced by poor weight gain, decreased bone mineral density, and reduced fertility (Bianco et al., 2007). On the other hand, TRPV5 null mice present less severe physiological consequences, nonetheless, the absence of TRPV5 channels causes hypercalciuria, compensatory hyperabsorption of dietary calcium, and abnormal bone thickness (Hoenderop et al., 2003). At negative membrane potentials, calcium-selective TRPV channels facilitate the passage of calcium ions to the cytoplasm. Calcium selectivity in TRPV5 and TRPV6 is accompanied by different mechanisms of Ca^2+^-dependent inactivation, helping to modulate Ca^2+^ entry during the first steps of transcellular transport (Hoenderop et al., 2001; Van Abel et al., 2005). One major difference between mammalian TRPV5 and TRPV6 channels rests in their distribution among tissues. Mammalian TRPV5 is predominantly expressed in the distal convoluted tubule and connecting tubule of the kidney, where likely to play a role in Ca^2+^ reabsorption. In contrast, mammalian TRPV6 is broadly expressed; it is found not only in the Ca^2+^ absorbing epithelia of the intestine but in placenta, pancreas and prostate, suggesting that the physiological roles of TRPV6 might not be limited to the maintenance of Ca^2+^ homeostasis (Peng et al., 2017).

Among vertebrates, it has been suggested that mammalian TRPV5 and TRPV6 channels originated from a duplication event in the last common ancestor of the group, between 312 and 177 million years ago (Peng, 2011; Saito et al., 2011). This would suggest that these genes belong to the mammalian clade exclusively and are not orthologs to TRPV5 and TRPV6 channels of other vertebrate groups. Besides these observations, not much else is known about the evolutionary history of these channels in vertebrates. Here we performed a comparative study of calcium-selective TRPV channels on representative species of all main groups of vertebrates. Thorough phylogenetic analyses in combination with functional assays allowed us to explore their evolutionary history and the evolution of functional features.

## Results

### Phylogenetic, primary sequence, and synteny analyses support the monophyly of the calcium-selective TRPV channels in vertebrates

To understand the duplicative history of calcium-selective TRPV channels, we first reconstructed a phylogenetic tree including sequences from species of all major groups of vertebrates. According to our maximum likelihood and Bayesian analyses, all node support measurements reached maximal values (100/1/100/1) for the monophyly of the group that includes all vertebrate calcium-selective TRPV channels (Figure 1). This indicates that all sequences included in our phylogenetic reconstruction derived from an ancestral sequence that was present in the genome of the vertebrate ancestor between 676 and 615 million years ago. A closer inspection of the sequence alignment also supports the monophyly of the group as several conserved residues can be easily mapped (Figure 1 - Figure supplement 1). Gaps and/or deletions in the alignment -introduced by the outgroups (TRPV1-4)- are not shared by any of the ingroup sequences (Figure 1; Figure 1 - figure supplement 1). The differences between the TRPV1-4 and TRPV5-6 clades can be traced to specific regions, including the ankyrin repeat domain (ARD), the Helix-loop-Helix domain (HLH), the intracellular linker between transmembrane segments 2 and 3 (S2-S3), and the pore domain (PD) (Figure 1 - Figure supplement 1). Moreover, synteny analyses also provide support for monophyly of the calcium-selective TRPV clade. Genes encoding TRPV5 and TRPV6 channels are located in a conserved genomic region, flanked by EPHB6 and KEL, suggesting that this genomic location was established early in vertebrate evolution and has remained relatively well conserved (Figure 1; Figure 1 - figure supplement 2). Thus, phylogenetic, primary sequence, and synteny analyses provide strong support for the monophyly of calcium-selective TRPV channels in vertebrates.

**Figure 1.**
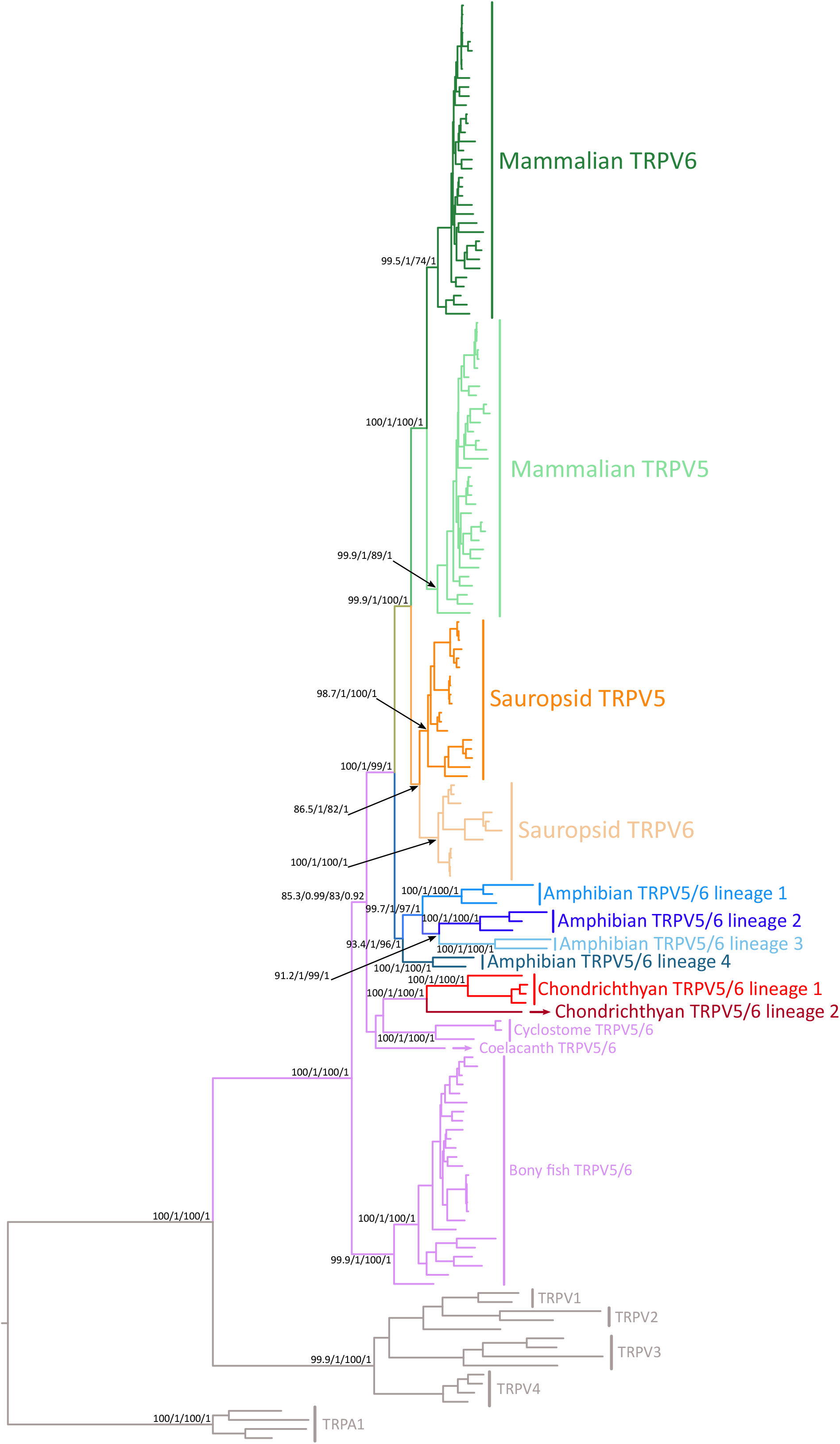
Maximum likelihood tree depicting evolutionary relationships among calcium-selective TRPV channels of vertebrates. Numbers above the nodes correspond to support from the Shimodaira-Hasegawa approximate likelihood-ratio test, aBayes, maximum likelihood ultrafast bootstrap and posterior probability values. TRPV1, TRPV2, TRPV3, TRPV4 and TRPA1 were used as outgroups. The scale denotes substitutions per site and colors represent gene lineages. The alignment used to build this phylogenetic tree, as well as, a phylogeny with species names for each terminal branch is found in the supplementary material section.

### Phylogenetic evidence for the independent origin of calcium-selective TRPV channels in mammals, sauropsids, amphibians, and chondrichthyans

Within the calcium-selective TRPV clade, our phylogenetic analyses suggest not a single but four independent expansion events during the evolutionary history of vertebrates (Figures 1 & 2). These can be noticed in our phylogenetic tree as duplicated copies in mammals, sauropsids, amphibians, and chondrichthyans are recovered sister to each other within each organismal group. For example, the sister group relationship between the mammalian TRPV5 and TRPV6 gene lineages is maximally supported (100/1/100/1) (Figure 1), reinforcing the hypothesis that the duplication event that gave rise to these copies occurred in the mammalian ancestor (Figure 2), as previously suggested (Peng, 2011; Saito et al., 2011). Similarly, the sister group relationship between the sauropsid gene lineages is also well supported (86.5/1/82/1) (Figure 1), indicating that a duplication event occurred in the sauropsid ancestor (Figure 2). Amphibians possess the most diverse repertoire of all vertebrates where four well-supported gene lineages were recovered (Figures 1 & 2). According to our phylogenetic tree, these amphibian gene lineages likely originate in the ancestor of the group (Figure 2). Such ancestor can be traced back to the carboniferous period, where amphibians were the dominant vertebrate terrestrial species (Wake and Koo, 2018). Similar to the situation described for mammals, sauropsids, and amphibians, our phylogenetic reconstruction supports a scenario in which the chondrichthyan (e.g. sharks, rays, and chimaeras) gene repertoire of calcium-selective TRPV channels was originated via gene duplication in the ancestor of the group (Figures 1 & 2). In contrast, the condition of a single gene copy (referred here as TRPV5/6) is present in cyclostomes, bony fish, and coelacanths (Figure 1). Thus, our results indicate that the gene repertoire observed in mammals, sauropsids, amphibians, and chondrichthyans originated independently in the ancestors of each group (Figure 2), suggesting that TRPV5 and TRPV6 in these groups are not 1:1 orthologs (Figure 2).

**Figure 2.**
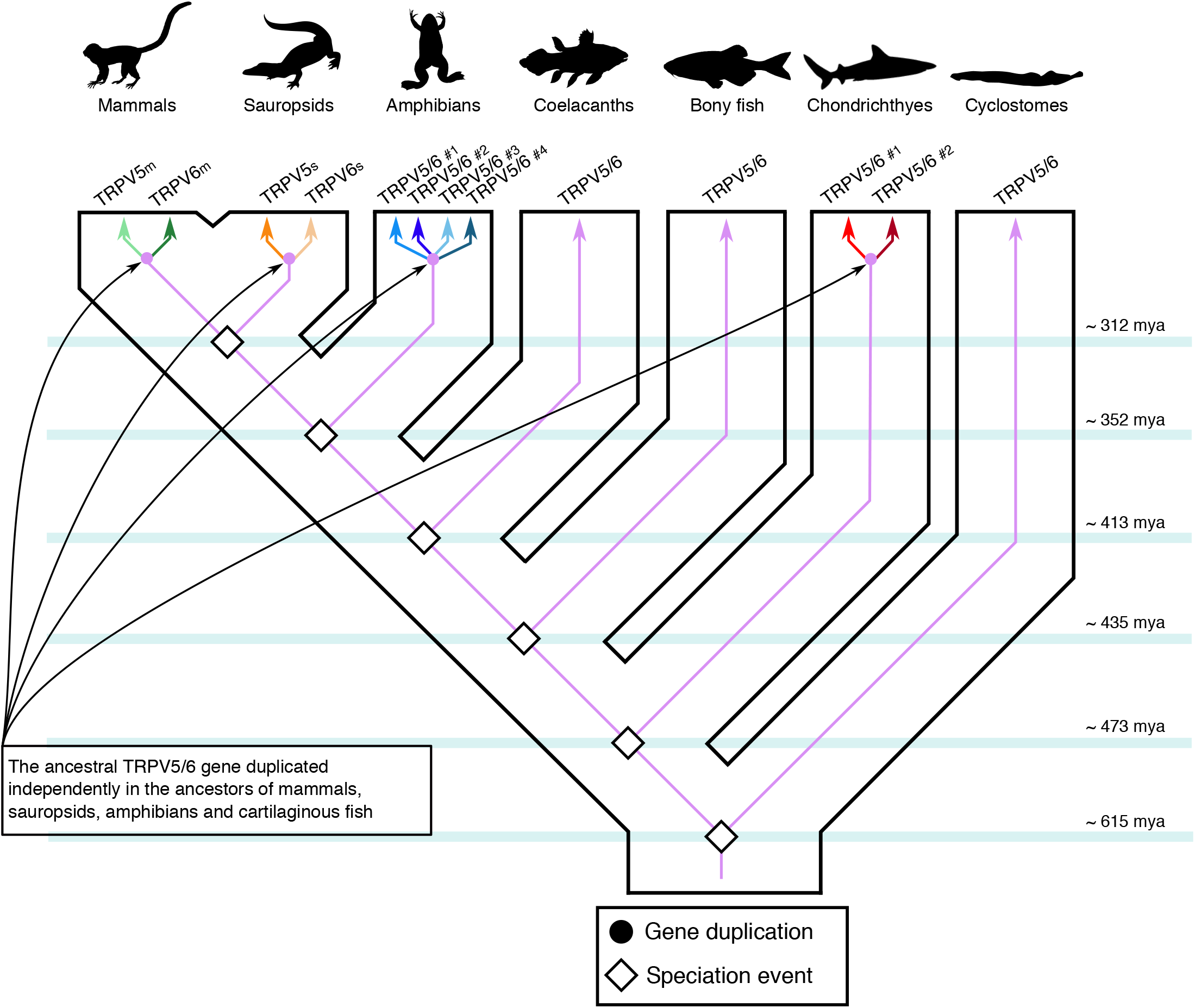
An evolutionary hypothesis regarding the evolution of the calcium-selective TRPV channels in vertebrates. According to our model the last common ancestor of vertebrates had a repertoire of one calcium-selective TRPV gene (TRPV5/6), a condition that has been maintained in cyclostomes, bony fish and coelacanths. In the ancestors of chondrichthyans, amphibians, sauropsids and mammals the single gene copy underwent independent duplication events giving rise to a more diverse repertoire of calcium-selective TRPV channels. In the case of chondrichthyans, sauropsids and mammals the single gene copy present in the ancestor of each group underwent a single duplication event-giving rise to a repertoire of two calcium-selective TRPV genes. In the case of amphibians, multiple duplication events in the ancestor of the group gave rise to a repertoire of four calcium-selective TRPV genes, for more details see figure 1. Vertebrate silhouette images were obtained from PhyloPic (http://phylopic.org/).

### Birds retained only one calcium-selective TRPV channel

When compared with other vertebrates, the case of sauropsids represents a special case of study. As in other tetrapods, the single gene copy present in the ancestor of the group underwent a duplication event originating a repertoire of two calcium-selective TRPV genes (Figures 1 & 3). However, during the radiation of the group, sauropsid TRPV5 was retained in all major lineages whereas sauropsid TRPV6 was found absent in birds (Figure 3). A comparison of the genomic regions in chicken and crocodile showed remaining exons of the sauropsid TRPV6 gene in the genome of chicken (Figure 3 - figure supplement 1). This suggests first that the gene present in birds corresponds to a sauropsid TRPV5 and not to a sauropsid TRPV6 as it has been previously suggested in the literature (Peng, 2011; Saito et al., 2011). The fact that birds can maintain calcium homeostasis according to their physiological requirements with just one of the calcium-selective TRPV channels could mean that a repertoire of two genes represent a case of functional redundancy in birds. In such case, the loss of one of the calcium-selective TRPV channels in the ancestor of birds could be understood as part of a stochastic process where the loss of the sauropsid TRPV6 is compensated by the retention of the sauropsid TRPV5 channel. Alternatively, it could be also possible that functional characteristics of the channel were deleterious for the avian physiology, in this case, the loss of TRPV6 channel might have been a product of purifying selection. With the available data it is not possible for us to distinguish between these two scenarios.

**Figure 3.**
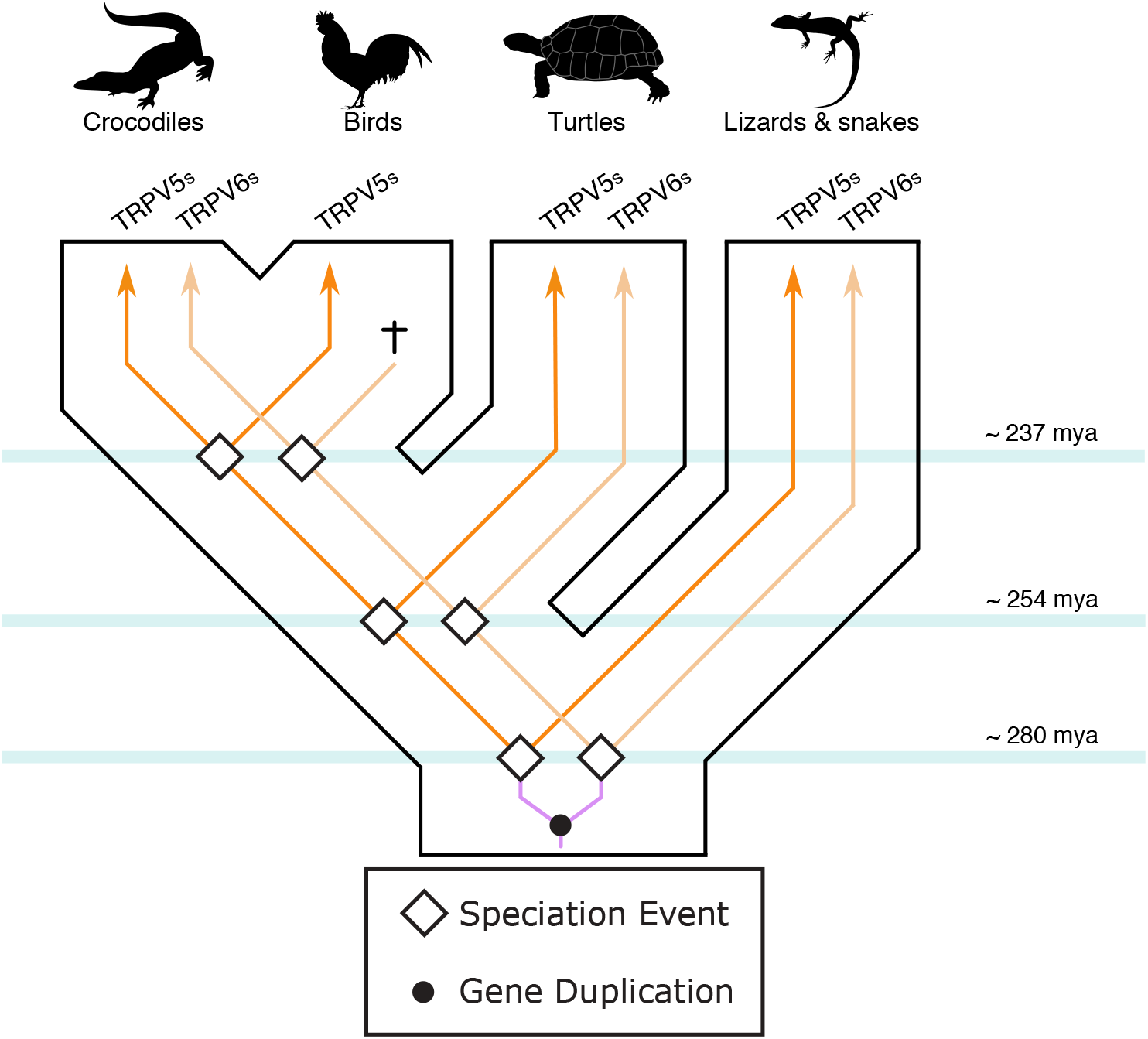
An evolutionary hypothesis regarding the evolution of the calcium-selective TRPV channels in sauropsids. According to this model the last common ancestor of sauropsids had a repertoire of one calcium-selective TRPV gene (TRPV5/6), which underwent a duplication event giving rise to a repertoire of two genes. During the radiation of the group the repertoire of two calcium-selective TRPV channels was maintained in crocodiles, turtles, lizards and snakes. However, in the ancestor of birds one of the genes was lost (sauropsid TRPV6), and the condition of a single gene copy was inherited by all descendant species. Vertebrate silhouette images were obtained from PhyloPic (http://phylopic.org/).

### Fast inactivation is a functional property associated to the duplication events

A closer inspection of the alignment used in our phylogenetic reconstruction revealed a remarkably strong signature sequence that correlates well with gene duplication events in all clades. This signature is composed of three amino acids located at the N-terminal helix-loop-helix (HLH) domain in mammals and sauropsids (Figure 4a). According to recently published structures (Hughes et al., 2018a; Singh et al., 2017), the HLH domain sits close to the S2-S3 intracellular linker and conveniently adjacent to the TRP domain helix (TD-helix), a known modulator of TRP channel gating (Gregorio-teruel et al., 2015; Valente et al., 2008) (Figure 4b). Previous studies have identified residues located in the S2-S3 linker (Nilius et al., 2002) and downstream of the transmembrane segment S6 (Suzuki et al., 2002) as structural modulators of inactivation in mammalian TRPV6 channels. Our sequence alignment also revealed a conserved pair of amino acids located in the S2-S3 linker. However, in contrast to the signature found at the HLH domain, the amino acids in the linker correlate with the duplication events only in mammals (Figure 4c). The presence of these signatures at two distant locations of the primary sequence, in channels that have duplicated independently, suggested to us a functional association between the S2-S3 linker, the HLH domain, and the TD-helix.

**Figure 4.**
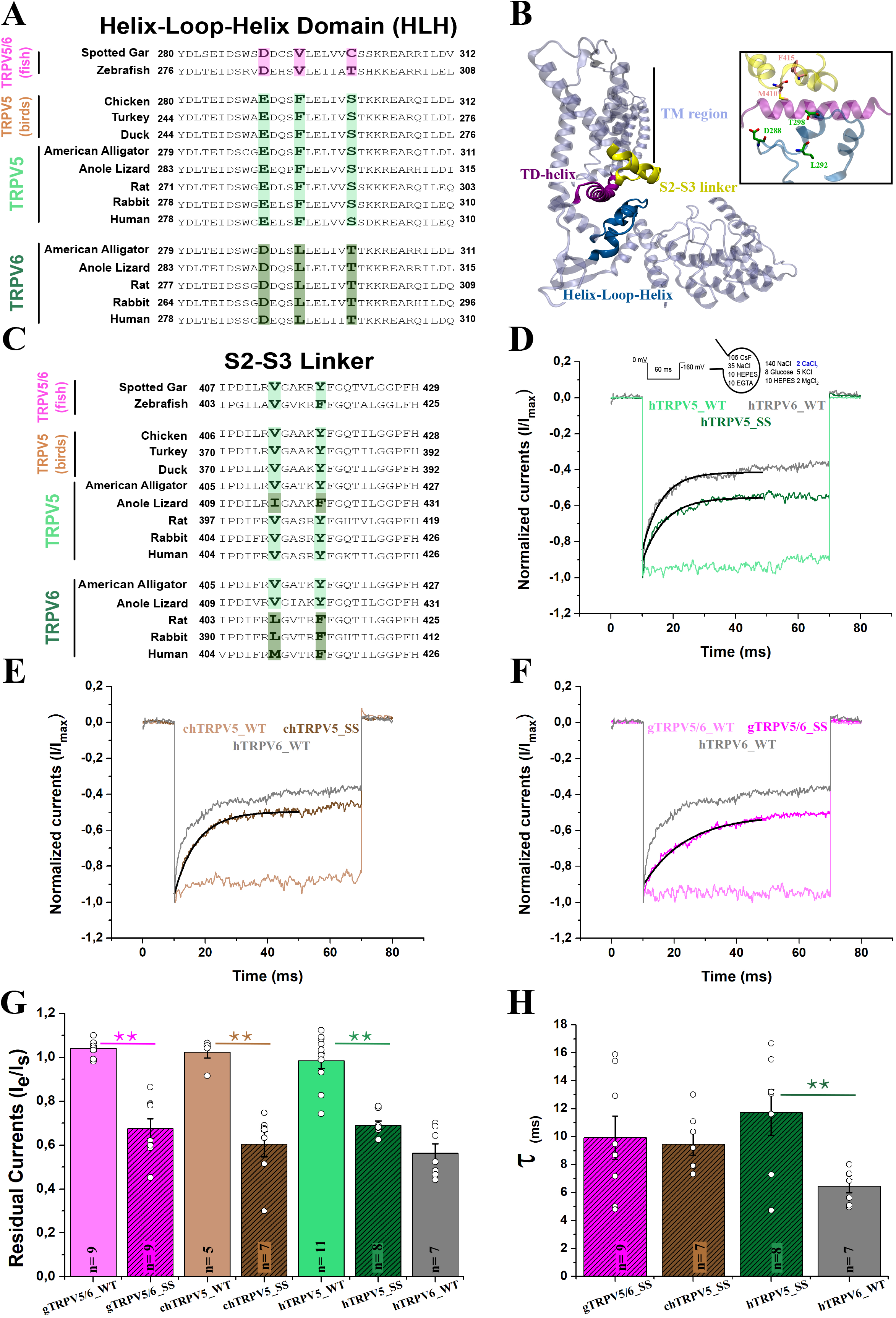
Specific amino acid replacements in the HLH domain and S2-S3 linker correlate with the fast inactivation phenotype. (A) Amino acid sequence alignment of the HLH domain. Conserved residues in TRPV5/6 (pink), TRPV5-type (light green) and TRPV6-type (dark green) channels are highlighted. (B) Structure of one subunit of the human TRPV6 channel (hTRPV6; PDB: 6E2F) (Singh et al., 2018) showing the structural triad HLH domain (blue), S2-S3 linker (yellow) and TD-helix (purple). TM region: Transmembrane region. The inset shows the location of the signature side chains in TRPV5. (C) Amino acid sequence alignment of the S2-S3 linker of the TRPV5-type, TRPV6-type and TRPV5/6 channels. Non-inactivating channels show conserved residues (light green) that differ from residues present in mammalian TRPV6 channels (dark green). (D) Normalized whole cell current traces recorded from transiently transfected HEK-293T cells expressing hTRPV5, hTRPV6, and engineered TRPV5 (hTRPV5_SS; E288D; F292L; S298T) containing the three-residue signature sequence at the HLH domain of TRPV6 channels. (E) Normalized whole cell current traces recorded from the chicken TRPV5 (chTRPV5_WT) and its triple mutant (chTRPV5_SS; E290D; F294L; S300T) containing the three-residue signature sequence at the HLH domain of TRPV6 channels and human TRPV6 channel (hTRPV6 WT). (F) Normalized whole cell current traces recorded from the spotted gar TRPV5/6 (gTRPV5/6_WT) and its mutant (chTRPV5/6_SS; V294L; C300T) containing the three-residue signature sequence at the HLH domain of TRPV6 channels and human TRPV6 channel (hTRPV6 WT). (G) Pooled data comparing the residual currents (see methods). (H) Pooled data comparing the inactivation time constants for inactivating clones (see methods). All current traces were recorded in the presence of 2 mM extracellular Ca^2+^ and in response to a voltage pulse at −160 mV for 60 ms from Vh=0 mV (Panels D; E; F). Black line indicates the fitting to a single exponential function (D; E; F). All bars represent mean value, errors represent S.E. and white dots represent values for each experiment (Panels G; H). ** represents p=0.01 and * p=0.05.

From a functional perspective, mammalian TRPV6 channels exhibit a characteristic two-component calcium-dependent inactivation (Hoenderop et al., 2001). In calcium-selective TRPVs, slow inactivation is determined by the binding of the Ca^2+^-Calmodulin complex to the C-terminal region of the channel and likely operates by the interaction of calmodulin’s C-lobe with the intracellular pore of the channel (Hughes et al., 2018b; Niemeyer et al., 2001; Nilius et al., 2003; Singh et al., 2018). Slow inactivation is present in both TRPV5 and TRPV6 channels from mammals. In contrast, fast inactivation depends solely on the presence of calcium ions and has been reported only in mammalian TRPV6 channels (Hoenderop et al., 2001; Nilius et al., 2002). Because of the role of the S2-S3 linker in fast inactivation (Nilius et al., 2002), we question whether these three amino acids at the HLH domain, represent not only a signature associated to duplication events but are also associated to a functional trait. To this end, we transferred the three amino acid signature found in the HLH domain of the human TRPV6 channel (hTRPV6) into the human TRPV5 channel (hTRPV5). Whole cell electrophysiological recordings showed that engineered hTRPV5 channels (hTRPV5_SS: E288D, F292L, S298T) now feature a robust fast inactivation in the presence of 2 mM extracellular [Ca^2+^] (Figure 4d and g). Wild type avian channels (*Gallus gallus*; chTRPV5), showed the absence of fast inactivation, as normally seen in mammalian TRPV5 channels (Figure 4e). In agreement with our previous results, engineered chTRPV5 channels (chTRPV5_SS: E290D, F294L, S300T) containing the amino acid signature from hTRPV6 now feature robust fast inactivation, mimicking mammalian TRPV6 channels (Figure 4e and g). Thus, our functional evidence shows that birds only kept a non-fast inactivating channel in their genomes. The absence of fast inactivation observed in the chicken TRPV5 channel does not rule out that other versions of the channel, likely similar to the ancestral condition, might exhibit fast inactivation. Thus, we also investigated inactivation in channels from vertebrates that possess the condition of single gene copy (e.g., spotted gar, *Lepisosteus oculatus*; gTRPV5/6). We observed that fast-inactivation is absent in gTRPV5/6 (Figure 4f), moreover -as seen in mammals and birds-fast inactivation can be easily introduced by mutagenesis at the HLH domain (gTRPV5/6_SS: V294L, C300T) (Figure 4f and g). Although all HLH mutants showed a similar time constant for fast inactivation (Figure 4h), it was somewhat slower than wild type hTRPV6. This suggests an additional element modulating inactivation, common to all and still not yet identified. Overall, our data not only highlights the importance of the HLH domain in fast inactivation but suggests that this phenotype represents an evolutionary innovation that originated with the expansion of the ion channel repertoire.

### Calcium-selective TRPVs from reptiles are intrinsically non-inactivating channels

Assuming a strict correlation between the duplication signature and the phenotype of fast inactivation, anole lizard’s TRPV5 channel (aTRPV5) should not fast-inactivate according to their sequence profile at the HLH domain (Figure 4a). In contrast, the residues at the S2-S3 linker (I415, F420) somewhat correspond to fast-inactivating mammalian TRPV6 channels (M410, F415) (Figure 4c). Electrophysiological recordings showed that the anole lizard’s TRPV5 channel elicits a modest fast inactivation where only 20% of the residual current is lost at steady state (Figure 5a). By introducing the sequence of non-inactivating channels at the S2-S3 linker (aTRPV5_VY: I415V, F420Y) we managed to completely abolish fast inactivation (Figure 5a). Moreover, the sole introduction of HLH signature in the anole lizard’s TRPV5 channel (aTRPV5_SS: E293D, F297L, S303T) produced a robust inactivation phenotype, similar to the one observed in the mammalian TRPV6 channel (Figure 5a and c). In contrast with the flexibility to adopt a new inactivation phenotype observed in all TRPV5-type channels analyzed (including aTRPV5), the anole lizard TRPV6 channel (aTRPV6) showed to be unsusceptible to inactivation. In this case, even the presence of both signatures shared by inactivating channels (aTRPV6_MF: HLH: D293, L297, T303; linker S2-S3: V415M, Y420F) was not enough to alter the non-inactivating nature of the channel (Figure 5c). Moreover, we found that the time constant of inactivation in both aTRPV5 and aTRPV5_SS (Figure 5d) was similar to other TRPV5-type mutants.

**Figure 5.**
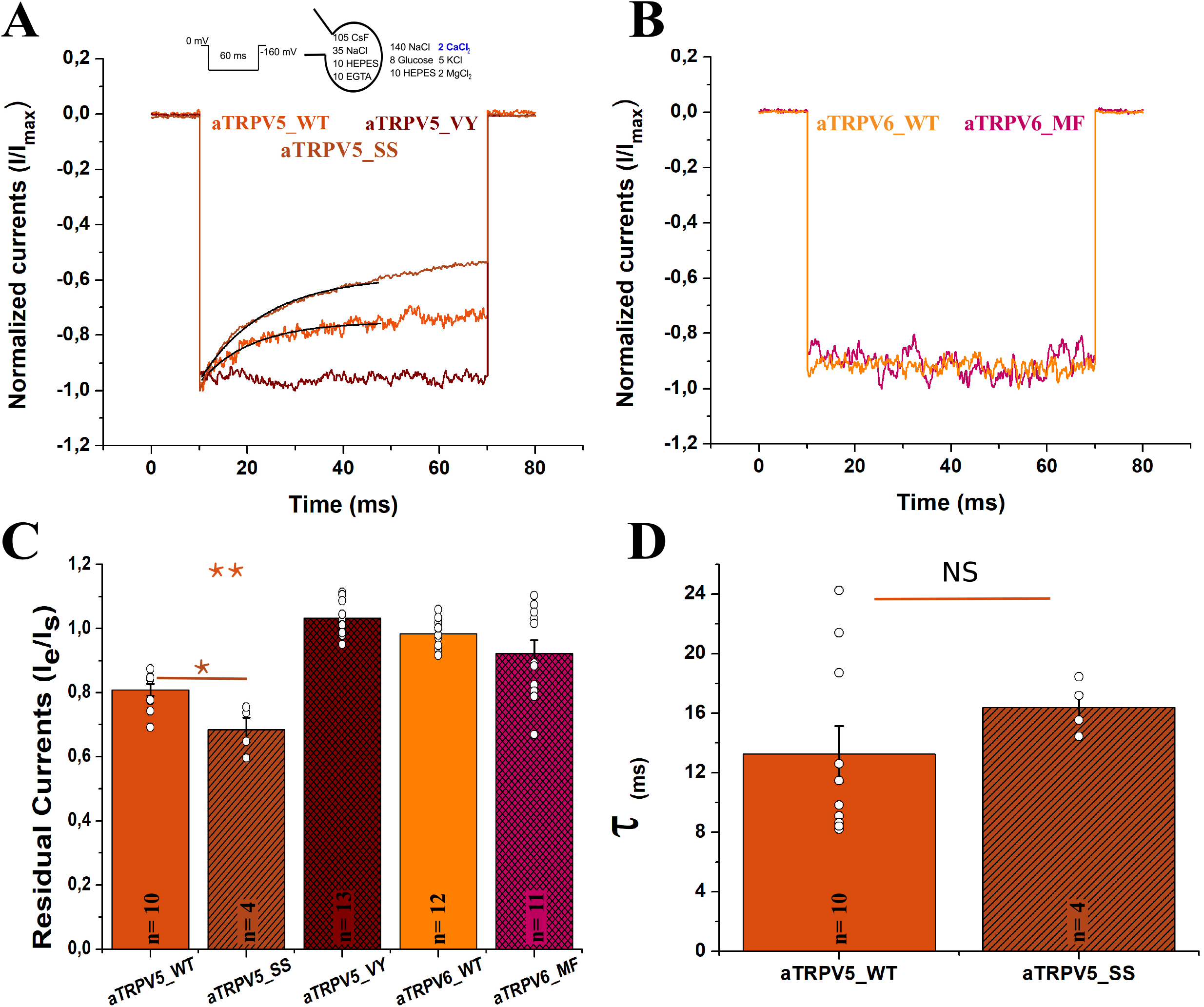
Calcium-selective TRPVs from reptiles lack a robust inactivation phenotype. (A) Normalized whole cell current traces recorded from transiently transfected HEK-293T cells expressing anole lizard TRPV5 channel wild type (aTRPV5 WT), containing the three-residue signature sequence of TRPV6 channels (aTRPV5_SS; E293D; F297L; S303T), and containing the amino acid signature corresponding to TRPV5 channels at the S2-S3 linker (aTRPV5_VY; I415V; F420Y). (B) Normalized whole cell current traces recorded from the anole lizard TRPV6 channel wild type (aTRPV6 WT) and containing the amino acid signature corresponding to human TRPV6 (aTRPV6_MF) (C) Pooled data comparing the residual currents. (D) Pooled data comparing the inactivation time constants for inactivating clones. All current traces were recorded in the presence of 2 mM extracellular Ca^2+^ and in response to a voltage pulse at −160 mV for 60 ms from Vh=0 mV (Panels A; B). Black line indicates the fitting to a single exponential function (A). All bars represent mean value, errors represent S.E. and white dots represent residual current value for each experiment (Panels C; D). ** represents p=0.01 and * p=0.05.

It has been reported that at high extracellular calcium (10 mM) both channels from human feature fast inactivation (Hoenderop et al., 2001). In our hands this holds true for calcium-selective TRPV channels from human, but absent in sauropsids (Figure 5 - figure supplement 1). These results suggest an additional modification in sauropsids, preventing inactivation by either modifying calcium binding or coupling. Such additional modification shouldn’t be present in the single copy gTRPV5/6, where high extracellular calcium readily induces fast inactivation (Figure 5 - figure supplement 1).

### Change of the expression profile from gills to kidneys in vertebrates

Based on expression profiles and inactivation properties it has been suggested that calcium-selective TRPV channels in mammals use a strategy in which the territories of expression are divided. While the mammalian TRPV6 channel should play a major role in Ca^2+^ absorption, mammalian TRPV5 would be mainly dedicated to calcium reabsorption in the kidney (Peng et al., 2017; Van Abel et al., 2005). This correlates well with their expression profiles; while mammalian TRPV6 channels are broadly expressed in a variety of tissues (including the kidney, intestine, placenta, pancreas, and prostate), mammalian TRPV5 is restricted to the distal convoluted tubule and connecting tubule in the kidney (Peng et al., 2017; Van Abel et al., 2005). In contrast to mammals, much less is known regarding the expression of these genes in other vertebrates. We then characterized the transcription profiles of calcium-selective TRPV channels in representative species of vertebrates (Figure 6).

**Figure 6.**
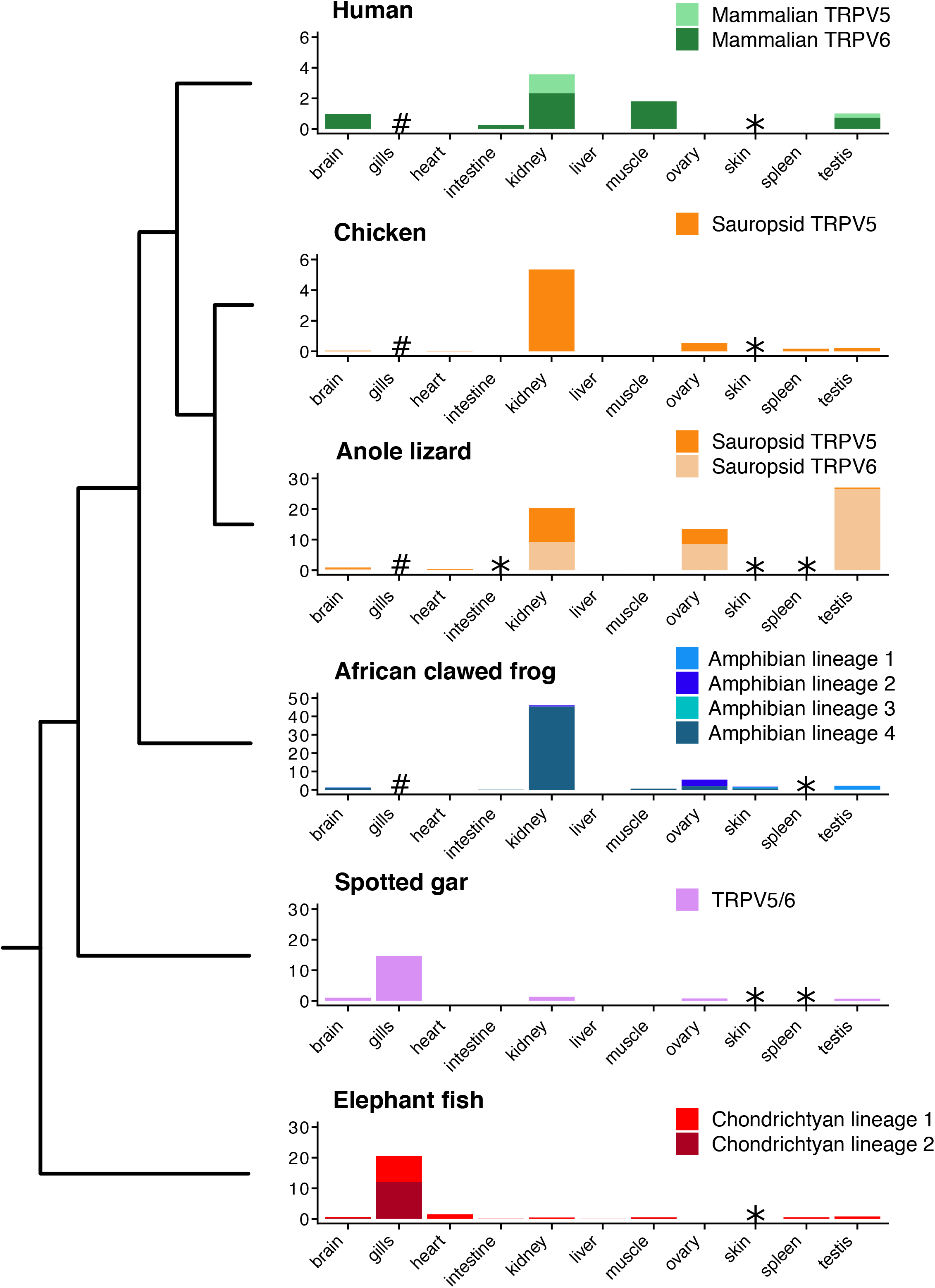
Comparative expression profiles of calcium-selective TRPV channels of vertebrate across multiple tissues. mRNA levels were estimated from public RNA-Seq libraries. Asterisks indicate data were not available for that species and tissue, and hashes indicate the tissue was absent. Gene expression was measured in transcript per million (TPM).

In agreement to what has been reported in the literature, our results show a localized expression for human TRPV5 channels while human TRPV6 is broadly expressed. Interestingly, both genes are well expressed in gonads, something that was found common to all surveyed species (Figure 6). In support of previous reports (Vanoevelen et al., 2011), we observed that aquatic vertebrates (i.e. spotted gar and elephant fish) mostly express calcium-selective TRPV channels in their gills, no matter the number of gene copies present in their genomes (Figure 6). The expression pattern observed in elephant fish suggests that the expansion of the gene repertoire did not give rise to a pattern in which the territories of expression were divided but rather served to increase the number of channels expressed in the gills (Figure 6). In contrast, the preferential expression of a single calcium-selective TRPV in kidney starts to be obvious in amphibians, where one of the four gene lineages present in their genome is clearly more abundant in this organ (Figure 6). The channels representing the other three gene lineages are present in other tissues such as skin, brain, and testis (Figure 6). This expansion in both gene repertoire and territories of expression might suggest early adaptations needed for terrestrial lifestyle (Mahasen, 2016). Thus, it seems that together with the conquest of land, the repertoire of genes not only expanded but also divided the territories of expression, while one of the copies is mainly expressed in kidneys, others are broadly expressed in a variety of tissues.

Sauropsids represent one more time an interesting case study. While birds retained only one of the two copies that were present in their ancestor, both genes are retained in all the other sauropsid groups (i.e. crocodiles, turtles, lizards, and snakes). In these groups, both gene lineages are expressed in the kidney and other tissues, while the sauropsid TRPV5 gene of birds is mainly expressed in the kidney and gonads (Figure 6). Thus, it seems that sauropsids have a more restricted expression of these genes and maximize calcium absorption by eliminating inactivation using different strategies. While reptiles might have been introducing structural modifications making the channel insensitive to calcium variations and/or becoming resistant to inactivation, birds just retained a non-inactivating channel.

## Discussion

In summary, our study shows that the evolutionary history of calcium-selective TRPV channels of vertebrates is mainly characterized by events of gene birth and death. According to our results, essential genes for calcium homeostasis have been duplicated four independent times during the evolutionary history of vertebrates. Our analysis suggests that among vertebrates, mammals, sauropsids, amphibians, and chondrichthyans, originated their repertoire of calcium-selective TRPV genes independently in the ancestors of each group. This observation implies that calcium-selective TRPV channels from these groups have different evolutionary origins. The independent origin of genes seems not to be uncommon in nature. Examples of these are ß-globin genes in vertebrates (Goodman et al., 1987; Hoffmann et al., 2018; Opazo et al., 2008), miRNA genes in monocots and brassicales (Gramzow et al., 2019), MHC class II genes in primates (Kriener et al., 2001), growth hormone genes in anthropoid primates (Li et al., 2005; Wallis and Wallis, 2002), growth differentiation factors (GDF1 and GDF3) in mammals and anurans (Opazo and Zavala, 2018), among others. In these cases, the function of gene products serves the same purpose, however, gene repertoires are not directly comparable as they do not have the same evolutionary origin. Thus, our observation of independent origin for the inactivation properties in calcium-selective TRPV channels opens an opportunity to understand in more detail the diversity of mechanisms associated with transepithelial Ca^2+^ transport and the evolution of the molecular coupling within the folded channel.

According to our data, it is likely that the ancestral condition of vertebrates corresponds to a single copy gene encoding a non-fast inactivating channel. This gene could have had properties similar to the ones observed in contemporary species of cyclostomes, bony fish, and coelacanths, including high selectivity for divalent ions (Qiu and Hogstrand, 2004). Tissue expression profiles highlight the relevance of the gills in calcium homeostasis in both sea and freshwater species. Among amniotes, duplicated copies are differentiated by specific amino acid substitutions in both the HLH domain and the S2-S3 linker. These amino acid substitutions could be strongly correlated with the phenotype of fast inactivation in all groups but reptiles, where additional modifications -outside of the region-analyzed here- might be relevant. The easiness by which the fast inactivation phenotype was introduced make us reason that an additional structural element could even predate the duplication events. Such element might not even be necessarily linked to inactivation originally. Under this scenario, fast inactivation could be a latent property that readily appeared multiple times in evolutionary history by just introducing specific mutations affecting the correct coupling between putative calcium binding site and the gate.

Amino acids at or close to the HLH domain have been implicated in the modulation of temperature-activation in TRPV1 and TRPV3 (Liu and Qin, 2017; Yao et al., 2011), and chemical-activation in TRPA1 (Saito et al., 2014). At the same time, the region close to the S2-S3 linker has been proposed as a calcium-binding site in TRPM4, TRPM2, and TRPC5 (Autzen et al., 2018; Duan et al., 2019; Huang et al., 2018), suggesting that calcium binding in the region may not be exclusive of TRPM channels. In calcium-selective TRPVs, the S2-S3 linker together with residues from the TD-helix and the inner gate have been associated to the different forms of calcium-dependent inactivation (Hughes et al., 2018b; Nilius et al., 2002; Singh et al., 2018; Suzuki et al., 2002). Therefore, the observations described here highlight the significance of the HLH motif - S2-S3 linker - TD-helix as a structural triad important for the modulation of TRP channel gating.

## Material and Methods

### Sequence data and phylogenetic analyses

We retrieved calcium-selective TRP channel sequences from representative species of all major groups of vertebrates. Our sampling included species from mammals, birds, reptiles, amphibians, coelacanths, holostean fish, teleost fish, chondrichthyans and cyclostomes (Supplementary File 1). Protein sequences were obtained from the Orthologous MAtrix project (OMA) (Altenhoff et al., 2018). In cases where the species were not included in the OMA project, we searched the NCBI database (refseq_genomes, htgs, and wgs) using tblastn (Altschul et al., 1990) with default settings. Protein sequences were aligned using the FFT-NS-1 strategy from MAFFT v.7 (Katoh and Standley, 2013). We used the proposed model tool of IQ-Tree v1.6.6 (Trifinopoulos et al., 2016) to select the best-fitting model of amino acid substitution (JTT + F + R6) (JTT: General matrix (Jones et al., 1992); F: empirical amino acid frequencies from the data; R6: FreeRate model with six categories (Soubrier et al., 2012; Yang, 1995). Phylogenetic relationships were estimated using maximum likelihood (ML) and Bayesian analyses. We employed maximum-likelihood to obtain the best tree using the program IQ-Tree v1.6.6 (Trifinopoulos et al., 2016). Support for the nodes was assessed with the Shimodaira-Hasegawa approximate likelihood-ratio test (SH-aLRT), the aBayes test from Anisimova et al., 2011 and 1,000 pseudoreplicates of the ultrafast bootstrap procedure (Hoang et al., 2017). Bayesian analyses were performed in MrBayes version 3.2 (Ronquist et al., 2012), running four simultaneous chains for 7 × 10^6^ generations, sampling trees every 2500 generations, and using default priors. We assessed convergence by measuring the standard deviation of the split frequency among parallel runs. Chains were considered to have converged once the average split frequency was lower than 0.01. We discarded trees collected before the chains reached convergence, and we summarized results with a majority-rule consensus of trees collected after convergence was reached. TRPV1, TRPV2, TRPV3, TRPV4 and TRPA1 were used as outgroups.

### Expression analysis

TRPV5 and TRPV6 expression was measured from a representative sample of vertebrates including elephant fish (*Callorhinchus milii)*, spotted gar (*Lepisosteus oculatus*), African clawed frog (*Xenopus laevis*), anole lizard (*Anolis carolinensis*), chicken (*Gallus gallus*) and human (*Homo sapiens*). RNASeq data from a spectrum of tissues including brain, heart, intestine, kidney, liver, muscle, ovary, testis, gills and skin, were gathered from the Short Read Archive (SRA). Accession numbers for species and tissue specific libraries can be found in Supplemental File 1. For all species except the elephant shark and the African clawed frog, we used Ensembl predicted gene sequences for gene expression references. Elephant fish and *X. laevis* cDNA sequences were collected from NCBI. To remove redundancy from libraries, all predicted TRPV5 and TRPV6 transcripts were removed from each library and replaced with our annotations. Adapter sequences were removed from the RNASeq reads using Trimmomatic 0.38 (Bolger et al., 2014), and reads were filtered for quality using the parameters HEADCROP:1, LEADING:30, TRAILING:30, SLIDINGWINDOW:5:30, and MINLEN:50. We used RSEM 1.3.1 (Li and Dewey, 2011) to map reads with Bowtie 1.2.2 (Langmead et al., 2009) and to estimate gene expression in units of TPM.

### Molecular Biology, Cell Culture, and Transfection

Open Reading Frames (ORF) encoding the different channels analyzed [Spotted gar TRPV5/6 WT (gTRPV5/6), gTRPV5 V294L C300T (SS), anole lizard TRPV5 WT (aTRPV5), aTRPV5 E293D F297L S303T (SS), aTRPV5 I415V F420Y (aTRPV5_VY), anole lizard TRPV6 WT (aTRPV6), aTRPV6 V415M Y420F (aTRPV6_MF), chicken TRPV5 WT (chTRPV5), chTRPV5 E290D F294L S300T (SS), human TRPV5 WT (hTRPV5), hTRPV5 E288D F292L S298T (SS) and human TRPV6 WT (hTRPV6)] were obtained from GenScript Biotech Corporation (Nanjing, China) inserted in a pcDNA3.1(+) vector. HEK 293T cells were grown in DMEM-F12 medium containing 10% (v/v) bovine fetal serum at 37°C in a humidity-controlled incubator with 5% (v/v) CO_2_. HEK 293T cells were transiently co-transfected with the different clones analyzed and peGFP-N1 to allow their identification.

### Electrophysiology and solutions

Whole-cell currents were measured with an Axopatch-200B amplifier. We used borosilicate pipettes (o.d. 1.5mm, i.d. 0.86mm, AM-Systems, Squim, WA) with resistance of 2 to 4.0 MΩ to form seals with access resistance between 2 and 6 GΩ. Macroscopic currents were recorded in response to a voltage step protocol from zero to −160 mV. Inactivation was analyzed in a time window of 60 ms. Recordings were digitized at 10 KHz and filtered at 5 KHz using a Digidata 1320 (Molecular Devices, LLC, California). The analysis was performed using Clampfit 10.3 (Molecular Devices, LLC, California). Fast inactivation was assessed by computing the residual currents, defined as the ratio of the current value at the end of the negative pulse over the current at the beginning of the voltage pulse (Derler et al., 2006). Inactivation time constants were obtained by fitting the current traces to a single exponential equation. The standard extracellular solution (Ringer-Na^+^) contained (in mM) 140 NaCl, 5 KCl, 2 CaCl_2_, 2 MgCl_2_, 8 glucose and 10 HEPES, at pH 7.4 adjusted with NaOH. Solution for high calcium extracellular concentration (~10 mM free calcium) contained (in mM): 125 NaCl, 10.5 CaCl_2_, 0.5 EGTA and 10mM HEPES, at pH 7.4 adjusted with NaOH. The standard internal (pipette) solution contained (in mM) 105 CsF, 35 NaCl, 10 EGTA and 10 HEPES, at pH 7.4 adjusted with CsOH. All experiments were performed at room temperature (20 to 25°C).

### Statistical Analysis

Data are expressed as mean ± S.E. Overall statistical significance was determined using analyses of variance (ANOVA one way) with a Bonferroni post-test, and T-student tests. For all conditions, the average was obtained from at least five independent experiments. Outliers were defined by using GraphPad QuickCalcs (https://www.graphpad.com/quickcalcs/Grubbs1.cfm), and removed from the analysis.

---

**Figure 1-figure supplement 1**. Amino acid sequence alignment of TRPV5 and TRPV6 channels. Human TRPV1-4 were used as outer groups. Species and genes are denoted to the left and amino acid numbers in the alignment at top. Insets show insertions introduced by TRPV1-4 sequences in the alignment. Relevant channel domains are highlighted on top: ARD: Ankyrin Repeat Domain; HLH: Helix-Loop-Helix Domain; S2-S3 linker: Intracellular loop between transmembrane segments 2 and 3; SF/p-Helix: Selectivity Filter & pore Helix; S6/TD-helix: Transmembrane Segment 6 & TRP Domain Helix.

**Figure 1-figure supplement 2**. Patterns of conserved synteny in the chromosomal regions that harbor the TRPV5 and TRPV6 genes. Asterisks indicate that the orientation of the genomic piece is from 3’ to 5’, gray lines represent genes that do not contribute to conserved synteny whereas diagonals indicate that the chromosomal pieces are not continuous. The colors of the TRPV5 and TRPV6 genes indicate the gene lineage to which they belong according to our phylogenetic tree shown in Figure 1.

**Figure 3-figure supplement 1**. Dot-plot of pairwise sequence similarity between the TRPV5 and TRPV6 genes of the American alligator (*Alligator mississippiensis*) and the corresponding syntenic region in the chicken (*Gallus gallus*). Light blue and light yellow vertical lines denote exons and introns, respectively. Vertical red lines denote exons that were lost from the chicken TRPV6 gene. Black lines indicate regions of high-quality alignment between the American alligator and chicken sequences, and the slope indicates the orientation. All exons of the TRPV5 gene are well conserved, however, for the TRPV6 gene there are several exons (red vertical lines) that do not align between the American alligator and chicken, being consistent with the loss of this gene in birds.

**Figure 5-figure supplement 1**. Comparison of fast inactivation phenotype at two different extracellular [Ca^2+^] concentrations. (A) Normalized whole cell current traces recorded from transiently transfected HEK-293T cells expressing spotted gar TRPV5/6, anole lizard TRPV5 (aTRPV5_WT), anole lizard TRPV6 (aTRPV6_WT), chicken TRPV5 (chTRPV5_WT), human TRPV5 (hTRPV5_WT), and human TRPV6 (hTRPV6_WT) (B) Pooled data comparing the residual currents. All current traces were recorded in response to a voltage pulse at −160 mV for 60 ms in the presence of 2 mM or 10 mM extracellular Ca^2+^. All bars represent mean value, errors represent S.E.M and white dots represent individual experiments. * represents p=0.05.

